# Single-Cell RNA Sequencing Reveals Commensal Microbes Amplify Sex-Specific Immune Programming in the Murine Lung

**DOI:** 10.64898/2026.01.22.700627

**Authors:** Alexandra Melton, Krista Ferrari, Joseph P Hoffmann, Kejing Song, Jay K Kolls, Janet E McCombs

## Abstract

Sex-based differences in respiratory disease outcomes are well recognized. However, the underlying immunological mechanisms driving this dimorphism remain incompletely understood. While sex hormones influence immune cell development and function, the role of commensal microbes in shaping sex-specific lung immunity has not been explored. Here, we used single-cell RNA sequencing (scRNAseq) and flow cytometry to profile lung immune cells in male and female mice housed under specific pathogen-free (SPF) or germ-free (GF) conditions. Under SPF conditions, males exhibited a striking myeloid bias, with increased monocytes and macrophages, along with broad upregulation of inflammatory mediators, including *S100a8, S100a9,* and *Il1b,* across multiple cell types, and enrichment of TNF and interferon (IFN) signaling pathways. In contrast, females displayed lymphocyte-skewed profiles, with higher frequencies of T cells and natural killer (NK) cells. Interestingly, these sex-based differences in immune composition and inflammatory programs were largely absent in GF mice, indicating that microbial exposure amplifies baseline immunological dimorphism between males and females. Notably, select sex-associated features, including female-biased NK cell enrichment, persisted irrespective of microbial status, suggesting intrinsic, microbiota-independent programming. Together, these findings indicate that commensal microbes modulate sex-specific lung immunity by amplifying pre-existing intrinsic differences, highlighting the intersection of extrinsic (microbial) and intrinsic (sex-linked) factors in shaping baseline mucosal immunity.

## Introduction

The lung is a uniquely regulated immune environment, constantly balancing the need for rapid pathogen defense with the risk of damaging inflammation. Sex differences in respiratory disease outcomes have long been recognized: males experience higher rates of severe disease and mortality from acute respiratory infections, whereas females typically mount more robust immune responses but show greater susceptibility to autoimmune disease^1–4^. These patterns reflect, in part, the effects of sex hormones and sex chromosome-linked immune regulators on tissue-resident and circulating immune cells^5–8^. While studies in peripheral blood and lymphoid organs have identified sex-specific patterns in certain immune cell subsets, whether these patterns hold in the lung itself remains unclear. Recent work in other mucosal tissues, such as the gut, has documented sex differences in immune cell populations. For example, female mice exhibit higher percentages of T cells in mesenteric lymph nodes ^9^. However, tissue-specific variation is likely, and the lung may follow distinct patterns. Direct hormonal regulation has also been demonstrated: testosterone increases alveolar macrophages and neutrophils while suppressing B cells in male lungs^10^. Even so, intrinsic factors alone may not fully account for the immune variation observed between males and females across mucosal tissues.

Commensal microbiota represent another major influence on immune homeostasis^11^. Often described in the context of the gut-lung axis, signals derived from the microbiota (including microbial metabolites, tonic pattern-recognition receptor (PRR) engagement, and baseline type I interferon (IFN) activity) shape systemic immune tone and, in turn, influence immune activation thresholds in the lung^12–14^. When these cues are disrupted, such as in germ-free or antibiotic-treated mice, antiviral defenses are weakened, myeloid responses become dysregulated, and inflammatory pathways are altered ^12,15^. Sex differences in microbiota composition have been observed in healthy mice and are associated with differences in baseline immune cell populations in intestinal-associated lymphoid tissues, such as the mesenteric lymph nodes^9^. These findings highlight the microbiota as a key environmental determinant of immune readiness and prompt investigation into how commensal microbes influence lung immunology at baseline.

Despite a growing body of sex-based studies on immunity, relatively little is known about how sex and microbial exposure interact at steady state, particularly within the adult lung. Most studies addressing sex differences in respiratory immunity have focused on infection models, leaving open the question of how these differences arise before an immune challenge occurs^1–3,7,16,17^. Whether commensal microbes amplify, constrain, or are required for sex-specific immune programs in the lung has not been directly explored.

To address this gap, we compared lung immune cell composition and transcriptional profiles in male and female adult mice raised under specific pathogen-free (SPF) or germ-free (GF) conditions using scRNAseq and flow cytometry. By examining sex and microbial status within the same experimental framework, our study provides a direct test of how intrinsic and environmental factors converge to shape baseline lung immunity. Our findings begin to shed light on the steady-state immune landscape that precedes, and likely contributes to, sex-specific differences in respiratory disease outcomes.

## Material and Methods

### Research Animals

For scRNAseq and 16S rRNA gene sequencing studies, six- to eight-week-old male and female germ-free (GF) and specific pathogen free (SPF) C57BL/6 mice were purchased from Taconic Biosciences. GF mice were shipped in sterile transport containers with food and water provided within the container. To preserve germ-free status, mice were maintained in their original shipping containers without transfer to new housing and were euthanized within three days of arrival.

For flow cytometry experiments, six- to eight-week-old male and female SPF C57BL/6 mice were purchased from Jackson Laboratories. Mice were housed in SPF conditions at Tulane University School of Medicine and provided food and water ad-libitum. All procedures were approved by Tulane University’s Institutional Animal Care and Use Committee and conducted in compliance with relevant ethical regulations for animal use.

### Immune Cell Isolation

Isolation of immune cells from the lungs of euthanized Taconic mice follows the protocol as described by Hoffmann et al with modifications where indicated^18^. In brief, both the left and right lungs of euthanized mice was collected in either sterile PBS and kept on ice for further processing. The media was removed, and the tissue was manually minced using sterile dissection scissors. Lung tissue was then resuspended in 2 mL IMDM (Gibco) containing 2 mg/mL collagenase type IV (Sigma-Aldrich) and 80 U/mL deoxyribonuclease (DNase) I (Sigma-Aldrich) and incubated with shaking at 233 rpm at 37°C for 1 hour. Following enzymatic digestion, the slurry was passed through a 70 µm cell strainer (Fisher) and red blood cells were removed using ACK lysis buffer (Gibco). Isolated cells were resuspended in IMDM containing 10% FBS (Hyclone) and counted using Nexcelom’s Cellometer Auto 2000 (Nexcelom) for downstream scRNAseq and flow cytometry.

Following euthanasia of Jackson labs mice, whole lungs were excised and placed on ice in 1 mL of IMDM with GlutaMAX (Gibco) supplemented with 1% antibiotic-antimycotic (Gibco). Lung samples were cut into ∼0.5 cm pieces and digested using 0.1 mg/mL DNase I (Roche) and 200 U/mL Collagenase Type IV (Worthington) for 45 minutes at 37℃ with gentle shaking every 15 minutes. Lung samples were sequentially passed through 70 µm and 40 µm cell strainers to generate a single-cell suspension. Cells were then washed in IMDM and 10% fetal bovine serum (FBS, Avantor) and centrifuged at 400 g for 5 minutes. Red blood cells were lysed using 1x ACK lysis buffer (Gibco) for 1.5 minutes. Following lysis, cells were washed and resuspended in IMDM containing 10% FBS. Viable cells were counted on a Nexcelom Cellometer prior to downstream flow cytometric staining and analysis.

### Single-Cell RNA Sequencing

Freshly isolated lung cells were processed for scRNAseq as described above for the Taconic mice. Single-cell RNA-seq libraries were generated using the 10x Genomics Chromium Fixed RNA Profiling workflow (Flex Gene Expression; Chromium Fixed RNA Profiling Reagent Kit) according to the manufacturer’s protocol. Briefly, viable single-cell suspensions were fixed using the crosslinking reagent provided in the kit and stored at - 80°C until processing. Target transcripts were captured using the Chromium Mouse Transcriptome Probe Set v1.0.1 (10x Genomics), and each sample was assigned a unique probe barcode. Following probe hybridization and washing, barcoded samples were pooled and loaded on 10x Chip Q to generate Gel Bead-in-Emulsions (GEMs). Within GEMs, probe pairs were ligated and extended to incorporate unique molecular identifiers (UMIs) and cell barcodes. Libraries were amplified by sample index PCR to add Illumina P5/P7 adapters and dual indexes. Library quality was assessed using an Agilent 2100 Bioanalyzer (High Sensitivity DNA kit) and quantified using a Qubit fluorometer. Libraries were sequenced at 750 pM on an Illumina NextSeq 2000 (P3 100-cycle flow cell; paired-end, dual-index). Raw sequencing data were processed using Cell Ranger version 7.1.0 (10x Genomics) and aligned to the mouse reference genome mm10-2020-A and the Chromium Mouse Transcriptome Probe Set v1.0.1.

### scRNAseq Analysis

Single-cell RNA sequencing (scRNAseq) data were processed and analyzed using the Seurat^19^ package (v5.2.1) in R. Individual Seurat objects were generated from the raw count matrices from each sample, which were then merged to generate a single Seurat object using the merge command. Cells with fewer than 200 or more than 5,000 detected genes were excluded to remove low-quality cells and potential doublets, while retaining highly transcriptionally active immune populations. Additionally, cells with >5% mitochondrial gene content were excluded to eliminate stressed or dying cells. Standard preprocessing steps included data normalization, scaling, dimensionality reduction, and graph-based clustering using a shared nearest neighbor approach. To correct for technical variation and inter-sample heterogeneity, batch correction was conducted using the Harmony integration algorithm^20^.

### Flow Cytometry

Single-cell suspensions from mouse lungs were prepared as described above and stained with fluorophore-conjugated antibodies (Supplemental Table 1). For Taconic mice samples, 1 x 10^6^ cells from each sample were added to the wells of a 96-well round bottom plate. Cells were washed with 3x with cold FACS buffer (1x PBS with 0.5% BSA). Blocking was achieved by resuspending cell pellets with 25µL FACS buffer containing CD16/CD32 rat anti mouse diluted 1:100 (v/v) and incubating for 30 minutes at 4°C. Following blocking, staining was achieved by adding 25 µL of fluorophore-conjugated antibody cocktail directly to each sample and incubating for 1 hour at 4°C. Cells were washed 3x, resuspended in FACS buffer, and analyzed using a Cytek Aurora spectral flow cytometer. Antibodies used for blocking and staining are as follows: Rat Anti-Mouse CD16/CD32 Fc Block (clone 2.4G2, BD Biosciences), APC Rat Anti-Mouse CD3e (clone 17A2, Biolegend).

For Jackson Laboratory samples, 1 × 10^^6^ cells were plated and incubated with Fc Block (Purified Rat Anti-Mouse CD16/CD32; BD Biosciences) for 30 minutes at room temperature in 1 PBS to block Fc receptors. Cells were then stained with Fixable Aqua Dead Cell Stain Kit (Invitrogen) and incubated for 30 minutes at room temperature, protected from light. Cells were washed in FACs buffer (1x PBS with 2% FBS) and then resuspended in 1x Stabilizing Fixative (BD Biosciences) for 20 minutes at room temperature, protected from light. After fixation, cells were washed and incubated overnight at 4°C in surface stain antibody cocktail containing Brilliant Stain Buffer (BD Bioscience), antibodies listed in Supplemental Table 1, and TrueStain Monocyte Block (BioLegend). Cells acquired on a Cytek Aurora 3-laser spectral flow cytometer (Cytek Biosciences). Data were analyzed in FlowJo v10 for Mac OS (BD Biosciences). Cells were subsequently gated to exclude debris and doublets prior to identification of CD45+ immune cells, with full gating strategies shown in Supplemental Fig. 2.

For the Taconic GF and SPF cohorts, a total of 20,000 events were acquired per sample; therefore, cell counts were derived directly from acquired events. For the Jackson Laboratory SPF cohort, total events varied across samples. To enable quantitative comparisons across samples, population counts were normalized to 10,000 live CD45+ single events using the DownSample plugin (FlowJo v10).

### 16S rRNA Gene Sequencing

Fecal pellets were collected as pooled samples from each of the four cages (one cage per group: GF females, GF males, SPF females, and SPF males) to verify microbial colonization status (SPF vs. GF). Because samples were pooled at the cage level and not collected from individual mice, this study does not address sex-specific differences in microbiota composition.

Genomic DNA was extracted from fecal samples using the Zymo Quick-DNA Fecal/Soil Microbe Kit (Zymo Research) following the manufacturer’s protocol. Bacterial community profiling was performed by 16S rRNA gene amplicon sequencing following Illumina’s MiSeq platform and the *16S Metagenomic Sequencing Library Preparation* workflow. Briefly, the V3-V4 regions of the bacterial 16S rRNA gene were amplified using region-specific primers with Illumina overhang adapters. Amplicons were purified, indexed, and pooled according to standard Illumina guidelines. Libraries were sequenced using a MiSeq v3 600-cycle kit (Illumina) with paired-end reads.

Raw reads were first trimmed to remove V3-V4 primer sequences using cutadapt ^21^(ran via command line). The resulting primer-trimmed reads were then processed with the DADA2^22^ pipeline in R for quality filtering, denoising, paired-end read merging, and chimera removal, resulting in inference of amplicon sequence variants (ASVs). Taxonomic assignment was performed using the SILVA^23,24^ ribosomal RNA gene reference database (version 138.1). A phyloseq^25^ object was created for downstream analysis, including taxonomic aggregation and calculation of alpha-diversity metrics (Observed richness, Shannon index, Simpson index). Relative abundance plots were generated at the genus level. Each sample represents a pooled cage-level fecal collection.

### Quantification and Statistical Analysis

Statistical analyses were performed using GraphPad Prism 10 for MacOS (GraphPad Software) and R version 4.5.1 (R Foundation for Statistical Computing). For flow cytometry analysis, unpaired Welch’s t-tests were used for two-group comparisons, and two-way ANOVA followed by Tukey’s multiple comparisons test was applied for analyses involving multiple groups or experimental conditions. Normality was assessed using the Shapiro-Wilk test where appropriate. Data are presented as mean +/- SEM unless otherwise indicated. Statistical significance was defined as p ≤ 0.05.

Differential gene expression (DEG) analysis was performed using the FindMarkers function in Seurat with the Wilcoxon rank-sum test and Bonferroni correction for multiple comparisons. Genes were considered differentially expressed based on an adjusted p-value ≤ 0.05 and an absolute log2 fold change (log2fc) ≥ 1.0. Gene set enrichment analysis (GSEA) was performed using ranked gene lists derived from all genes tested in each comparison, ordered by log2fc. Enrichment of Hallmark^26^ biological pathways (The Broad Institute Molecular Signature Database^27^ (MSigDB)) was assessed using normalized enrichment scores (NES), with pathways considered significant based on a false discovery rate (FDR) < 0.1.

## Results

### Sex-based differences in lung immune cell composition in SPF mice

To characterize baseline sex differences in lung immune cell composition, we performed single-cell RNA sequencing (scRNAseq) on cells isolated from the lungs of age-matched male and female SPF mice (**Fig. 1A, Supplemental Fig. 1A**). UMAP clustering identified 27 transcriptionally distinct clusters corresponding to 10 major cell populations, including T cells, B cells, NK cells, monocytes (Monos), alveolar and interstitial macrophages (Macs), endothelial cells (Endo), epithelial cells, fibroblasts (Fibro), and proliferating cells (**Figs. 1B, 1C**). Although overall clustering patterns were similar between sexes, immune cell proportions differed markedly. Females exhibited a higher frequency of lymphocytes, comprising 79-83% of lung cells compared with 54-58% in males, driven largely by elevated T and NK cell proportions. In contrast, male lungs were enriched in myeloid cells, with threefold higher frequencies than females (37-39% vs. 12-16%), primarily due to expanded monocytes and interstitial macrophages (**Fig. 1D, 1E**).

**Fig 1.**
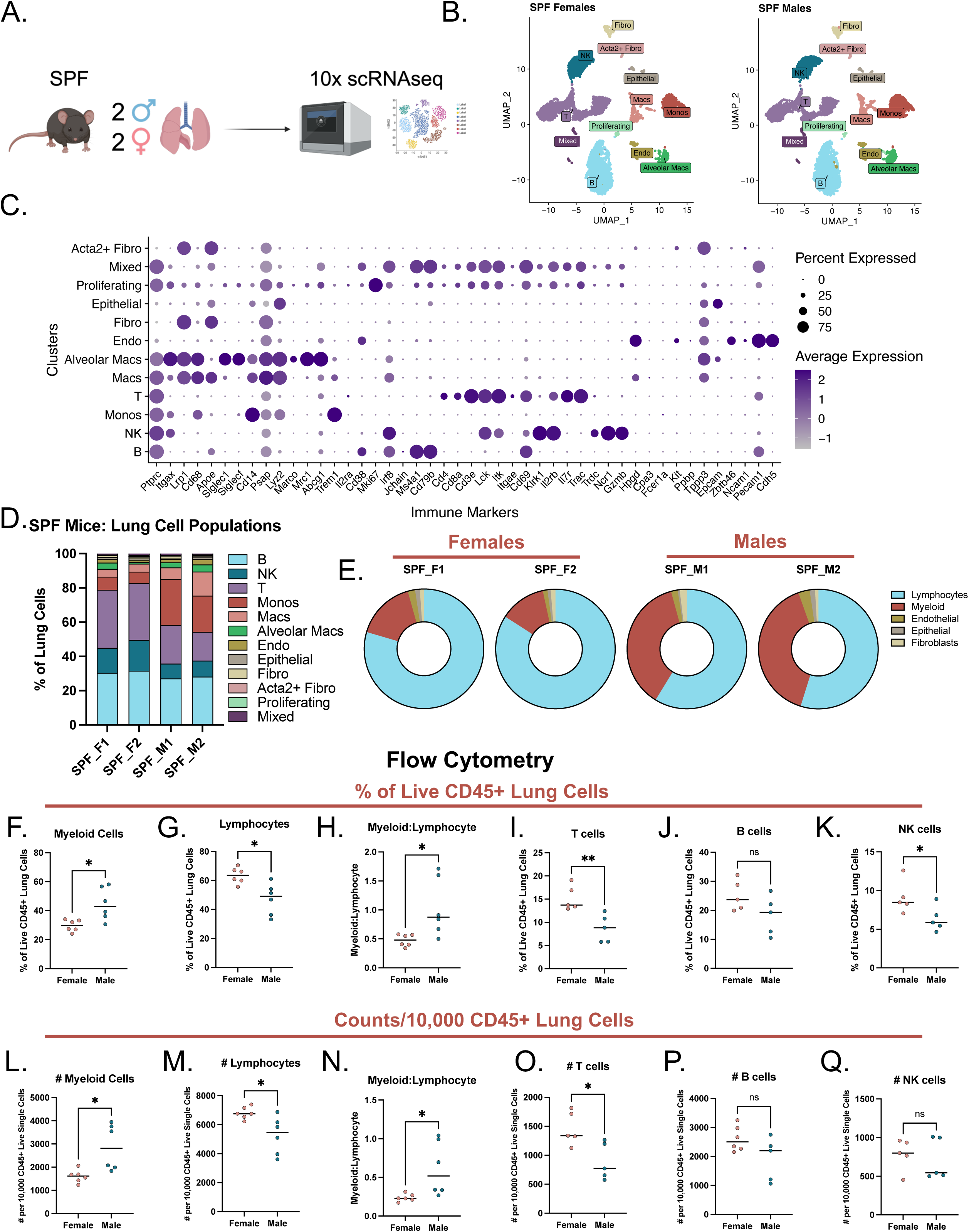
Single-cell RNA sequencing (scRNAseq) and flow cytometry analysis of lung cells isolated from specific pathogen-free (SPF) male and female mice. A. Study design. scRNAseq was performed on lung cells harvested from naïve male (M) and female (F) SPF mice. Created using BioRender (https://BioRender.com) B. UMAP plots illustrating annotated cell populations. C. Dot plot of gene markers used for cell type identification. D. Proportions of major lung cell populations in individual female (F1/F2) and male (M1/M2) mice. E. Pie charts illustrating broad cell categories. F-Q. Flow cytometry of lung cells isolated from SPF male and female mice. F-G. Proportions of myeloid (F) and lymphoid (G) cells. H. Myeloid to lymphocyte ratio. I-K. Lymphocyte subsets. Proportions of T (I), B (J), and NK (K) cells. L-Q. Counts of indicated populations normalized to 10,000 Live CD45⁺ Single Cells (DownSample plugin, FlowJo). F-Q. Unpaired Welch’s t-tests used to determine significance (*p < 0.05, **p< 0.01). Fibro = fibroblasts, Endo = endothelial, Mono = monocytes, Macs = macrophages. A-E: n = 2 per group. F-K: n=5 or 6 per group.

Flow cytometry analysis confirmed these observations, revealing significantly higher frequencies of myeloid cells in male lungs compared to females, resulting in a markedly elevated myeloid-to-lymphocyte ratio (mean: 1.03 vs. 0.48, respectively; **Figs. 1F-1H** & **Supplemental Fig. 2A**). Notably, this pattern remained consistent with the scRNAseq results despite the use of mice from different vendors (Jackson Laboratories for flow cytometry vs. Taconic for scRNAseq), supporting the robustness of these sex-specific findings. Female lungs harbored significantly higher proportions of T cells and NK cells, and while B cells trended higher in females, this did not reach statistical significance (**Figs. 1I-1K**). Analysis of cell numbers revealed a similar pattern; males had significantly more myeloid cells, whereas females displayed significantly higher lymphocyte counts, with T cells driving this difference (**Figs. 1L-1O**). B cell and NK cell counts did not differ significantly between sexes, although both trended higher in females (**Figs. 1P-1Q**).

### Sex-specific transcriptional programs in SPF lung immune cells

To assess the influence of sex on transcriptional programs we then performed differential gene expression (DEG) analysis. We identified 320 genes upregulated in males and 60 in females (log₂FC ≥ 1.0, adjusted p ≤ 0.05), with 14,008 genes shared between groups (**Fig. 2A**). The male-enriched gene signature was characterized by proinflammatory and myeloid-associated genes, including alarmins^28^ (*S100a8, S100a9*), acute phase protein^29^ *Lcn2*, proinflammatory mediators^30,31^ (*Il1b, Cxcl2, Ccl6*), and pattern recognition molecules (*Cd14*) (**Fig. 2B, 2C**). In contrast, *B2m*, which encodes a key structural component of major histocompatibility complex class I molecules (MHC I), was the top DEG upregulated in females. Given its broad expression across cell types (**Fig. 2C, Supplemental Fig. 3A**), elevated baseline *B2m* expression may reflect enhanced MHCI-mediated antigen presentation capacity, potentially contributing to the more robust CD8⁺ T cell response to vaccination and infection observed in females^5,32–34^. Gene set enrichment analysis (GSEA) of Hallmark pathways revealed that male lung cells were significantly enriched for inflammatory signaling pathways, including TNF via NF-κB, IL-6/STAT3, and interferon responses (**Fig. 2D**). Conversely, female lung cells were enriched for cell cycle-related pathways (E2F and MYC targets, and G2M checkpoint, DNA repair), suggesting higher baseline proliferative activity in females.

**Fig 2.**
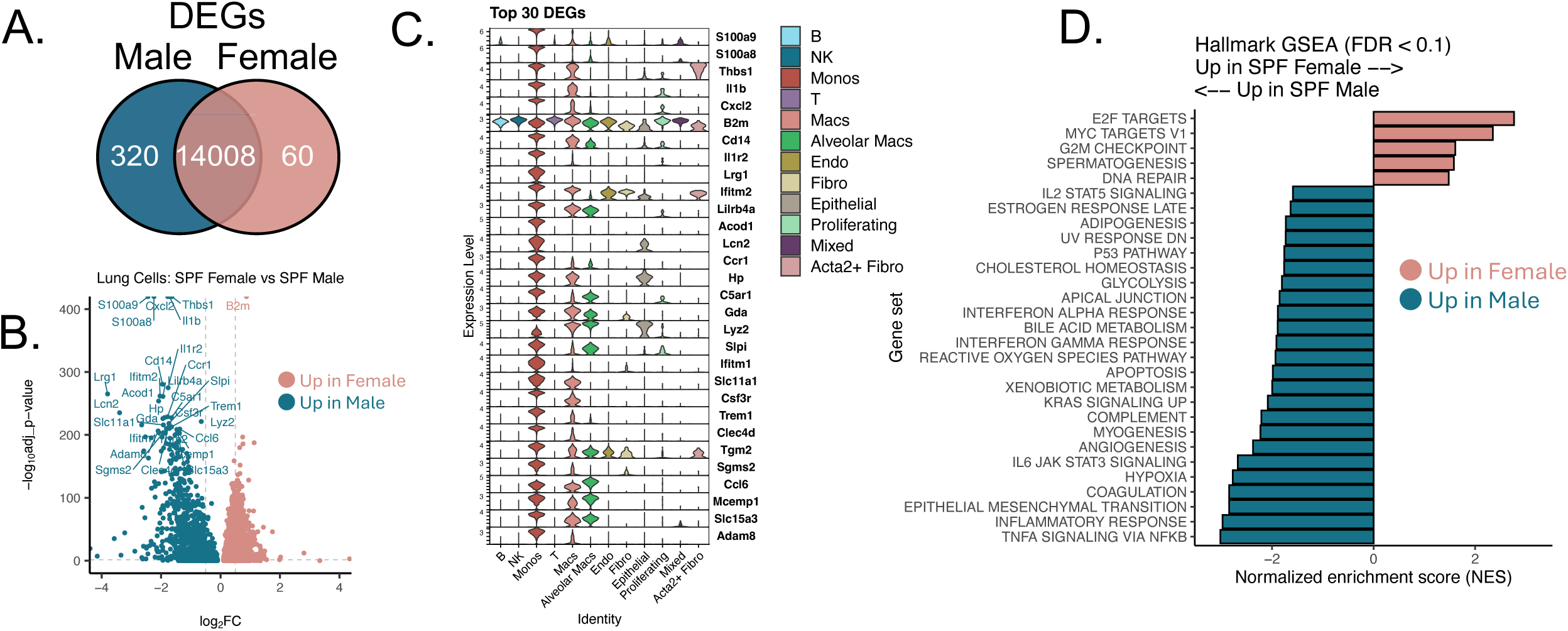
Differentially expressed genes (DEG) analysis comparing lung cells isolated from specific pathogen-free (SPF) male and female mice. **A.** Venn diagrams depicting the number of DEGs identified by comparing lung cells from male (teal) and female (peach) SPF mice. **B.** Volcano plot highlighting top 30 DEGs (log2fc ≥ 1.0 or ≤ -1.0, adjusted p-value ≤ 0.05). **C.** Distribution of top 30 DEGs across cell types. **D.** Normalized net enrichment scores (NES) from gene set enrichment analysis (GSEA) of Hallmark biological pathways. Significance determined using a false discovery rate (FDR) cutoff of 0.1. (All genes were used in GSEA. Gene list was ranked by abs(log2fc)). Fibro = fibroblasts, Endo = endothelial, Mono = monocytes, Macs = macrophages.

### Microbiota amplifies sex-based immune differences in the lung

Our baseline SPF analyses revealed striking sex-based differences in lung immune cell composition: males with elevated myeloid cell frequencies, and females with a more lymphocyte-biased profile. To examine whether these sex-specific immune differences persist in the absence of commensal microbes, we performed scRNAseq on lung cells isolated from germ-free (GF) male and female mice (**Fig. 3A**). To verify microbial status, we performed 16S rRNA sequencing on fecal samples from the SPF and GF mice (**Supplemental Fig. 1B**). Unsurprisingly, DNA from GF samples fell below the threshold for reliable sequencing, consistent with successful maintenance of germ-free conditions. SPF mice harbored relatively diverse bacterial communities, typical of murine gut microbiota^9,35^ (**Supplemental Fig. 1B**). Alpha diversity metrics revealed similar richness and evenness between SPF males and females, indicating no significant sex-based differences in baseline microbial composition (**Supplemental Fig. 1C**). For scRNAseq, we applied the same preprocessing, clustering, and annotation methods as used for the SPF analysis (**Figs. 1B, 1C**). As seen with the SPF dataset, overall clustering patterns were similar between male and female GF mice (**Fig. 3B**). Strikingly, however, the pronounced sex-based differences in lymphocyte and myeloid proportions observed in SPF mice were largely absent in GF mice (**Fig. 3C, 3D**).

**Fig 3.**
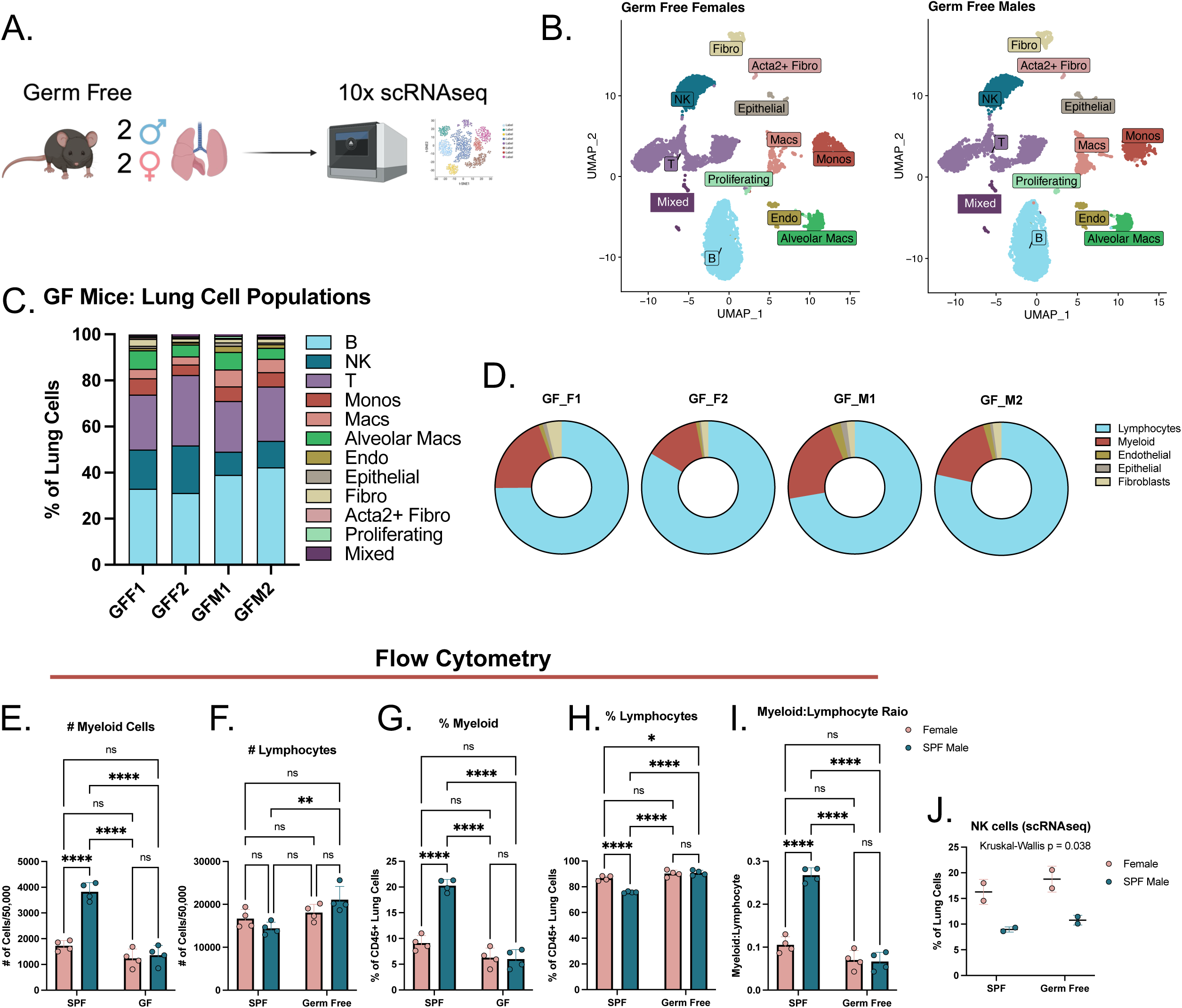
Single-cell RNA sequencing (scRNAseq) and flow cytometry analysis of lung cells isolated from germ-free (GF) and specific pathogen-free (SPF) male and female mice. **A.** Study design. scRNAseq was performed on lung cells harvested from naïve male and female GF mice. Figure created using BioRender (https://BioRender.com) **B.** UMAP plots illustrating annotated cell populations. **C.** Proportions of lung cell populations in individual female (F1/F2) and male (M1/M2) GF mice. **D.** Donut plots summarizing broad cell type composition. **E-I.** Flow cytometry analysis of lung cells illustrating myeloid and lymphocyte cell counts (**E,F**), proportions (**G,H**) and the myeloid to lymphocyte ratio (**I**) across GF and SPF conditions. **J.** NK cell proportions in lungs of SPF and GF male and female mice identified by scRNAseq. For panels A-D & J, n = 2 per group. Panels E-I, Bar graphs show mean ± SEM for each group. Two-way ANOVA and Tukey’s multiple comparisons test used to determine significance (Shapiro-Wilk for normality). *p < 0.05, **p<0.01, ***p<0.001, ****p<0.0001), n = 4 per group. Fibro = fibroblasts, Endo = endothelial, Mono = monocytes, Macs = macrophages

Flow cytometry revealed similar results. Male SPF mice had significantly higher myeloid cell frequencies and myeloid-to-lymphocyte ratios compared to both SPF and GF females (**Figs. 3E-3I** & **Supplemental Fig. 2B**). In contrast, GF male and female mice had comparable myeloid cell proportions and no significant difference in the myeloid-to-lymphocyte ratio. Notably, scRNAseq revealed higher proportions of NK cells in both SPF and GF female Taconic mice (**Fig. 3J).** This female bias was confirmed by flow cytometry in SPF Jackson mice, in which NK cell frequencies, but not absolute counts, were significantly higher in females (**Fig. 1Q**).

### Differential gene expression in GF and SPF lung immune cells

To examine the impact of commensal microbial exposure on gene expression, we performed differential gene expression analysis comparing lung immune cells from GF and SPF mice. Overall, 164 genes were upregulated in SPF mice compared to 78 in GF mice, with 14,562 genes shared between the two groups (**Fig. 4A**). Interestingly, when we examined which sex drove these SPF-enriched genes, 155 were predominantly upregulated in males and only 6 in females, with 3 genes elevated in both sexes. In contrast, GF mice showed minimal sex bias, with only 23 of the 78 GF-enriched genes elevated in males, 26 in females, and 29 shared between the sexes. Cell type specific analysis revealed that monocytes (138 DEGs) and macrophages (97 DEGs) accounted for the largest transcriptional changes in males, while T cells contributed most to the differences in females (39 DEGs) (**Fig. 4B**; **Supplemental Fig. 1D**). Notably, we observed 380 DEGs between male and female SPF mice (**Fig. 2A**) compared to 242 DEGs between SPF and GF mice (**Fig. 4A**), suggesting that sex exerts a greater influence than microbial status on transcriptional variation in lung immune cells, though microbiota exposure appears to amplify sex-based differences.

**Fig 4.**
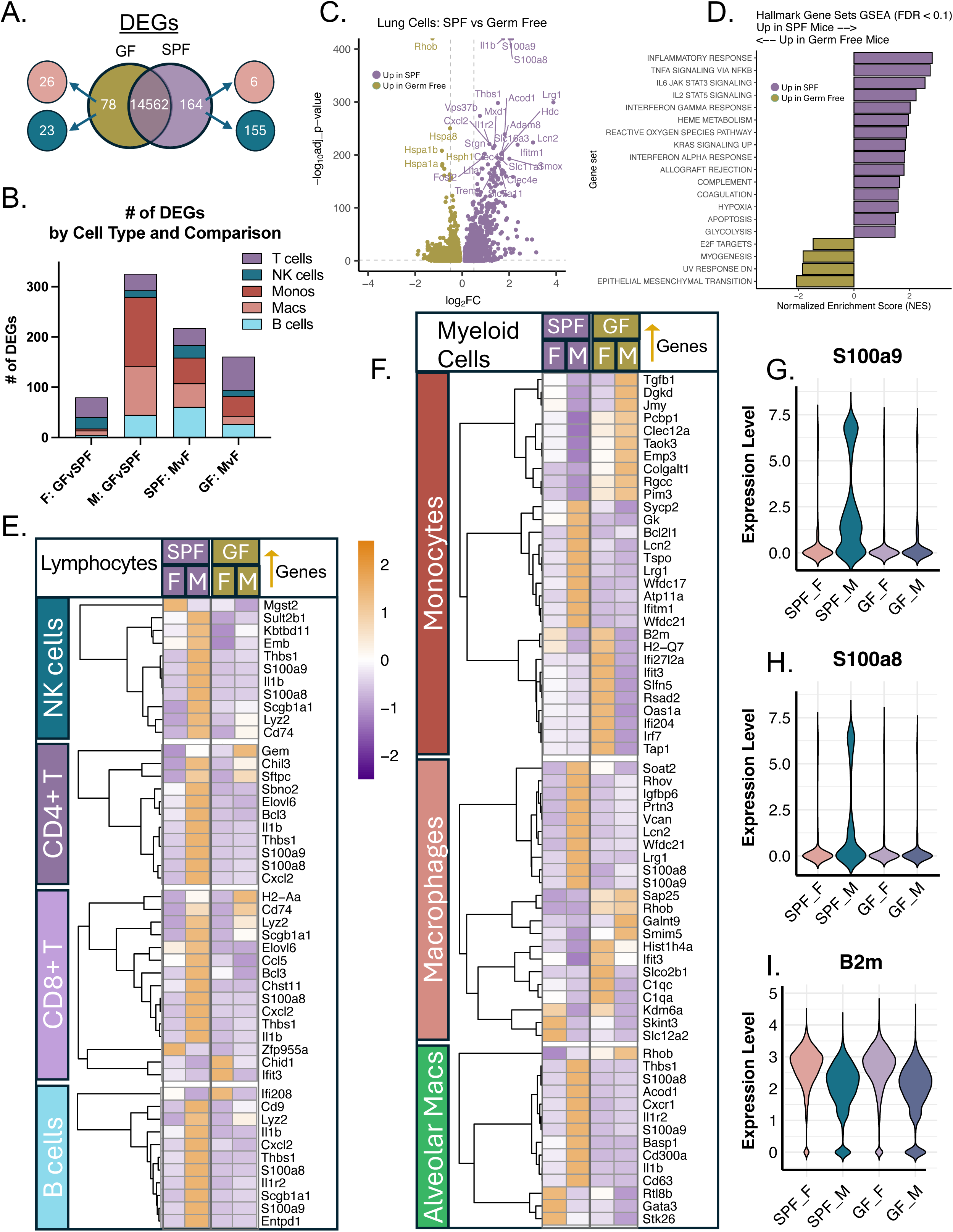
Differentially expressed genes (DEG) analysis comparing lung cells isolated from germ-free (GF) and specific pathogen-free (SPF) male and female mice. **A.** Venn diagrams of DEG overlap between GF (gold) and SPF (purple) mice. Of the DEGs identified, the number of genes up in females (peach) and males (teal) is also displayed. **B.** Number of DEGs per cell type and comparison group (F = Female, M = Male). DEG count based on the number of significantly upregulated or downregulated genes (|log₂FC| ≥ 1.0, adjusted p ≤ 0.05) per cell type (B cells, macrophages (Macs), monocytes (Monos), natural killer (NK) cells, and T cells). **C.** Volcano plot of top DEGs between SPF and GF mice. Top 30 genes up in SPF (purple) or GF (gold) are indicated. **D.** Hallmark gene set enrichment analysis (GSEA) of DEGs. Pathways with FDR < 0.1 shown with normalized enrichment scores (NES). **E-F.** Heatmap illustrating the top 10 upregulated DEGs per condition for myeloid **(E)** and lymphocyte **(F)** populations. Rows were hierarchically clustered by Pearson correlation of scaled expression; columns are ordered by condition and sex. Values represent z-scored average gene expression within each cell type. (adjusted *p* < 0.05, log₂ fold change > 1, expressed in ≥ 10% of cells per group). n = 2 per group.

Transcripts enriched in SPF lung immune cells included inflammatory mediators such as *S100a8*, *S100a9*, *Il1b*, *Cxcl2*, and *Thbs1*, whereas those enriched in GF lungs were primarily associated with stress responses, including heat shock proteins (*Hspa8*, *Hspa1b*, *Hspa1a*, and *Hsph1*) (**Fig. 4C**). GSEA further demonstrated that inflammatory pathways, including TNF signaling, IL-6/STAT3, interferon responses, and complement activation, were enriched in SPF mice (**Fig. 4D**). In contrast, few pathways were enriched in GF mice consistent with a lower baseline inflammatory state.

To examine cell type-specific differences in gene expression, we performed hierarchical clustering of the top DEGs across cell types (**Figs. 4E-4F**). In myeloid cells, SPF males exhibited pronounced upregulation of inflammatory genes compared to GF males and both female groups (**Fig. 4F**). Lymphocyte populations also displayed sex- and microbiota-dependent expression patterns, though the magnitude of differences was more moderate than in myeloid cells (**Fig. 4E).** Notably, several proinflammatory genes, including *S100a8, S100a9,* and *Il1b*, were consistently upregulated across multiple myeloid and lymphoid cell types in SPF males, suggesting a coordinated inflammatory program rather than cell-type specific responses (**Figs. 4E-4H, Supplemental Fig. 3A**). Interestingly, we found that *B2m* was also highly expressed in GF females across all cell types, similar to the SPF females, suggesting intrinsic programming independent of microbiota (**Fig. 4I, Supplemental Fig. 3A**).

## Discussion

Sex-based differences in lung immunity are well recognized across respiratory infections and inflammatory diseases, yet the underlying drivers of these baseline differences remain incompletely understood. Intrinsic factors, such as sex hormones and sex chromosome-linked immune regulators, influence lung-resident immune populations^5,10^, but how these intrinsic influences intersect with environmental cues, particularly microbial exposure, to shape baseline immune diversity is unclear. Using scRNAseq and flow cytometry to compare SPF and GF mice, we found that sex-based differences in lung immune composition and transcriptional activity were pronounced under SPF conditions. Males showed myeloid cell enrichment and heightened inflammatory gene signatures, whereas females displayed a lymphocyte-biased profile. Strikingly, these differences were largely absent in GF mice. We also noted elevated NK cell proportions in females that persisted regardless of microbial status, suggesting intrinsic sex-based programming. Together, these findings suggest that commensal microbiota amplify sex-based immune diversity in the lung, highlighting a key intersection between intrinsic and environmental factors in shaping baseline lung immunology.

Commensal microbes influence lung immunity through several mechanisms. Microbial metabolites, tonic stimulation of PRRs, and baseline type I IFN signaling establish the steady-state activation threshold of innate and adaptive immune cells^12–14,36,37^. These pathways modulate myeloid cell maturation, neutrophil responsiveness, dendritic cell activation thresholds, and basal inflammatory tone ^12–14,36,37^. Studies using germ-free or antibiotic-treated mice have demonstrated impaired antiviral responses, weakened type I IFN priming, and altered Th17/Treg balance, underscoring the role of commensals in maintaining mucosal immune readiness^12,15^. These mechanisms provide a plausible explanation for why sex differences were robust in SPF mice but collapsed under GF conditions in our study.

While sex differences in immune cell populations have been documented in other mucosal tissues, such as the gut, where females exhibit higher percentages of T cells in mesenteric lymph nodes, relatively little is known about how sex and microbial exposure interact specifically in the lung. Most studies addressing sex differences in immunity have focused on systemic compartments or other mucosal sites, and tissue-specific variation in these patterns is likely. Our data complement prior work demonstrating hormonal regulation of tissue-specific immunity. Testosterone has been shown to increase alveolar macrophages and neutrophils while also suppressing B cells in male lungs, consistent with the elevated myeloid cell frequencies we observed in the SPF males in our study^10^. Building on this, our data reveal a dimension not captured by hormonal studies alone and suggest that commensal microbial exposure amplifies or may be required for these sex-specific differences to fully manifest in the lung.

Here, we found that SPF males showed a strong myeloid bias, characterized by increased monocytes and macrophages and marked upregulation of inflammatory mediators, including *S100a8*, *S100a9*, *Il1b*, and *Cxcl2.* These alarmins and chemotactic factors promote neutrophil recruitment and amplify innate immune responses, and have been linked to severe inflammatory lung injury ^28,38,39^. Gene set enrichment analysis further revealed enrichment of TNF, IL-6/STAT3, and interferon signaling in males, pathways associated with heightened disease severity during respiratory infections^3^. In contrast, SPF females had greater frequencies of T cells and NK cells along with enrichment of cell cycle-related pathways. Though our study examined steady-state conditions, these baseline differences may help explain clinical observations in which males exhibit greater inflammatory pathology during respiratory infections^2,3,40–42^, whereas females often generate more robust lymphocyte-mediated responses^5,16,17^.

The pronounced myeloid-to-lymphocyte differences observed between male and female SPF mice were almost entirely absent in GF mice. And though select sex-linked transcriptional features were detectable in germ-free mice, their impact on immune cell composition and inflammatory gene expression was markedly attenuated in the absence of commensal microbes. In addition to this, differential gene expression analysis comparing SPF and GF mice showed that SPF-associated transcriptional changes were dominated by myeloid populations in males, whereas females exhibited more modest, T cell-driven differences. These findings suggest that microbial exposure differentially influences innate and adaptive immunity in males and females. This microbiota-hormone interaction may help explain why sex differences in respiratory disease susceptibility often emerge or intensify after puberty, when both hormonal and microbial factors converge to shape tissue-resident immunity^8,43–45^.

Although microbial status strongly influenced immune cell composition in the lung, sex accounted for a greater number of differentially expressed genes overall, highlighting the dominance of intrinsic sex-linked transcriptional programs. One example of this was the increased NK cell frequencies observed in females across both SPF and GF conditions. This persistence in GF mice suggests that NK cell programming in the lung is less dependent on microbial cues than are the myeloid inflammatory pathways. Given the critical role of NK cells in early innate immune responses to viral infection, higher baseline NK frequencies in female lungs may enhance early immune surveillance and antiviral responsiveness, potentially contributing to more efficient viral control^46^.

Notably, *S100a8* and *S100a9* were among the most consistently upregulated genes in SPF males, revealing a coordinated inflammatory signature that extended beyond classical myeloid cells. While expression was highest in monocytes, *S100a8/a9* was also elevated across lymphocytes, epithelial cells, endothelial cells, and fibroblasts. *S100a8/a9* encode calcium-binding proteins that form the calprotectin complex, a potent alarmin that amplifies inflammation through TLR4 and RAGE signaling^47^. In humans, elevated S100A8/A9 expression marks severe inflammatory lung disease and is especially prominent in airway epithelium during COVID-19^39^. The broad induction of *S100a8/a9* across immune and epithelial compartments in SPF males suggests that commensal microbes prime basal activation of this inflammatory axis, potentially predisposing males to dysregulated inflammatory responses often observed during severe pulmonary infections, such as those caused by SARS-CoV-2.

An interesting gene upregulated in both SPF and GF females was *B2m*. *B2m* encodes β2-microglobulin, a critical structural component of MHC class I molecules essential for antigen presentation to CD8^+^ T cells^48^. Elevated *B2m* expression across multiple cell types in females may enhance baseline antigen presentation capacity, potentially contributing to the more robust CD8^+^ T cell responses to vaccination and infection observed in females ^5,32–34^. This heightened antigen presentation capacity may also provide a mechanistic link to the increased prevalence of autoimmune diseases in females^49^, a hypothesis that warrants further investigation.

## Conclusion

Our findings suggest that sex and microbiota do not act independently but rather interact to shape baseline lung immunity. We found that commensal microbes amplify sex-specific immune programming in the lung, driving inflammatory, myeloid-skewed profiles in males and lymphocyte-biased states in females. Understanding this interplay may guide the development of targeted interventions that account for both microbial and hormonal influences on respiratory health.

## Study Limitations

This study focused on baseline immune differences in the absence of infection. Therefore, the functional consequences of the observed sex- and microbiota-dependent differences remain to be determined. While key findings were validated by flow cytometry, our conclusions are largely based on transcriptomic data, which may not fully reflect protein expression or cellular function. Due to cost constraints, our scRNAseq analysis was performed using two mice per group. Future studies with larger sample sizes will be needed to confirm our findings. While sex differences in microbiota composition have been documented in mice^9^, our study used 16S sequencing solely to verify SPF vs. GF status using cage-level pooled fecal samples. This approach precluded analysis of individual-level sex-specific microbial profiles, and future studies with individual-level microbiota profiling would help identify whether specific microbial taxa or metabolites drive the sex-dependent lung immune phenotypes we observed. Additionally, germ-free models have developmental and physiological differences beyond microbial exposure that may influence immune profiles^50^. Finally, we focused on young adult mice (6-8 weeks old); thus, age-related changes in sex-specific immune programming will require future investigation.

## Acknowledgements

We thank C. Porretta (Tulane University School of Medicine) for assistance with flow cytometry experiments and Lori Rowe and Clara Krzykwa of the Molecular Virology and Sequencing Core (RRID: SCR_027144), at Tulane National Biomedical Research Center (P51 Base Grant-P5110D011104 RRID: SCR_008167) for performing 16S sequencing. Research reported in this publication was supported by the National Institute of General Medical Sciences of the National Institutes of Health under Award Number P20GM152305. The content is solely the responsibility of the authors and does not necessarily represent the official views of the National Institutes of Health.

## Figures

**Supplemental Fig 1. Additional data supporting Figures 1 and 3 A.** Dot plot showing the expression of selected sex-linked genes across specific pathogen-free (SPF) and germ-free (GF) male and female lung samples. Male mice exhibit high expression of Y-linked genes (Uty, Ddx3y, Eif2s3y, and Kdm5d), while females show high expression of X chromosome genes (Eif2s3x, Kdm6a, and Kdm5c). **B.** Top 10 most abundant genera detected in fecal samples from SPF male and female mice. Fecal pellets were collected at the cage level, and microbial composition was determined by 16S rRNA gene sequencing. Bars represent relative abundance of the top 10 genera, with all remaining taxa grouped as “Other”. **C.** Phylum-level alpha diversity of the fecal microbiome in SPF mice. Each point represents a pooled cage sample. Diversity metrics shown include observed richness (number of phyla), Shannon index (richness and evenness), and Simpson index (dominance). Each point represents a pooled fecal sample from each group.

**Supplemental Fig 2. Additional data supporting Figures 1 and 3. A and B.** Flow cytometry gating strategies and representative plots corresponding to **Figures 1F-1Q (A) and 3E-3I (B)**. **C.** Heatmap displaying the number of DEGs across major immune cell types (B cells, macrophages (Macs), monocytes (Monos), natural killer (NK) cells, and T cells) for each comparison shown in **Figure 4B**. DEG count was based on the number of significantly upregulated or downregulated genes (|log₂FC| ≥ 1.0, adjusted p ≤ 0.05). Log2-transformed DEG counts were used for visualization.

**Supplemental Fig 3. Additional data supporting Figure 4. A.** Violin plots illustrating expression of S100a8/9 and B2m across major cell types.

## Abbreviations

SPF: specific pathogen free
GF: germ free
scRNAseq: single-cell RNA sequencing
MHC I: major histocompatibility complex class I
IFN: Interferon
PPR: pattern-recognition receptor
DEG: differentially expressed genes
GSEA: Gene set enrichment analysis
MSigDB: Molecular Signature Database
NES: normalized enrichment score
FDR: false discovery rate
log2fc: log2 fold change
GEO: Gene Expression Omnibus
UMAP: Uniform Manifold Approximation and Projection
Monos: monocytes
Macs: macrophages
NK: natural killer
Endo: endothelial cells
Fibro: fibroblasts

