## Supplemental Figures 1-3 for "Single-Cell RNA Sequencing Reveals Commensal Microbes Amplify Sex-Specific Immune Programming in the Murine Lung"

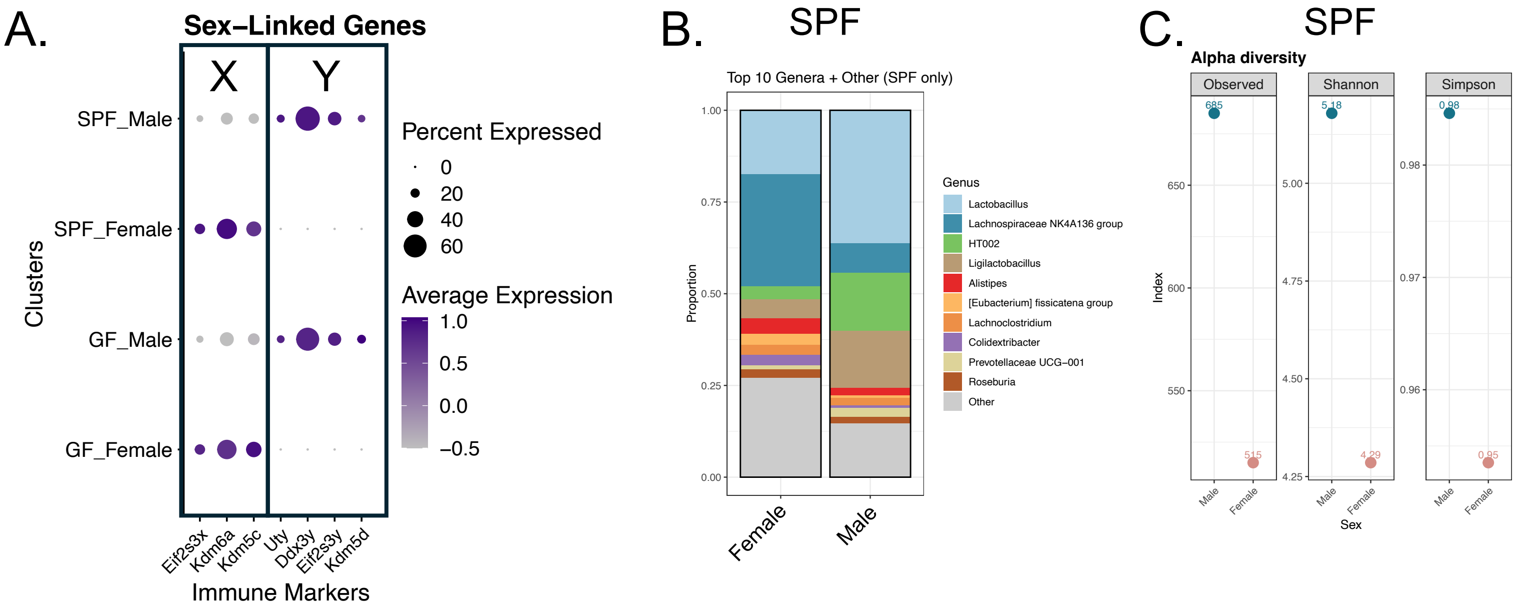


**Supplemental Fig 1. Additional data supporting Figures 1 and 3 A.** Dot plot showing the expression of selected sex-linked genes across specific pathogen-free (SPF) and germ-free (GF) male and female lung samples. Male mice exhibit high expression of Y-linked genes (Uty, Ddx3y, Eif2s3y, and Kdm5d), while females show high expression of X chromosome genes (Eif2s3x, Kdm6a, and Kdm5c). **B.** Top 10 most abundant genera detected in fecal samples from SPF male and female mice. Fecal pellets were collected at the cage level, and microbial composition was determined by 16S rRNA gene sequencing. Bars represent relative abundance of the top 10 genera, with all remaining taxa grouped as "Other". **C.** Phylum-level alpha diversity of the fecal microbiome in SPF mice. Each point represents a pooled cage sample. Diversity metrics shown include observed richness (number of phyla), Shannon index (richness and evenness), and Simpson index (dominance). Each point represents a pooled fecal sample from each group.

### Supplemental

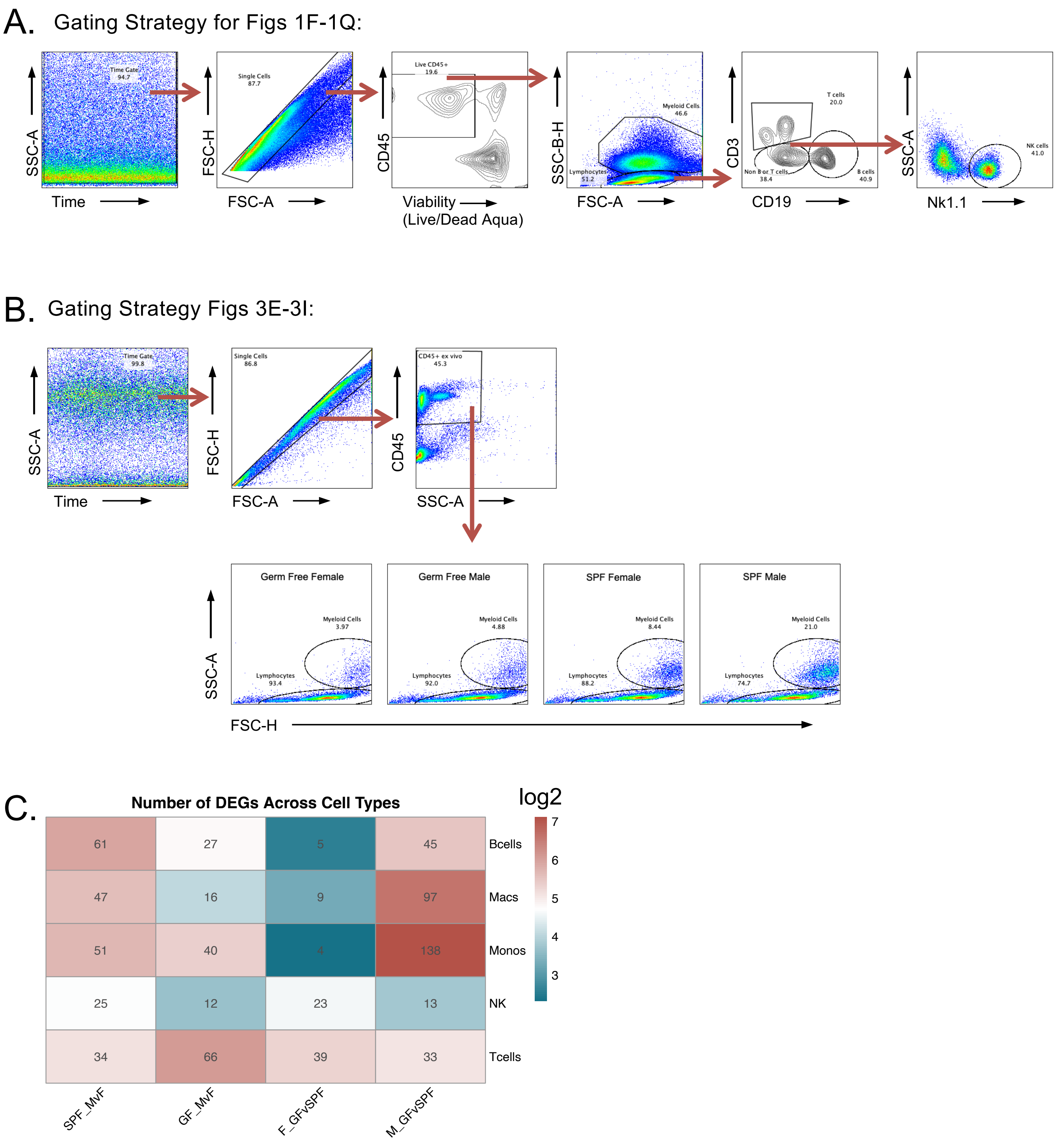


**Supplemental Fig 2. Additional data supporting Figures 1 and 3. A and B.** Flow cytometry gating strategies and representative plots corresponding to **Figures 1F-1Q (A) and 3E-3I (B)**. **C.** Heatmap displaying the number of DEGs across major immune cell types (B cells, macrophages (Macs), monocytes (Monos), natural killer (NK) cells, and T cells) for each comparison shown in **Figure 4B**. DEG count was based on the number of significantly upregulated or downregulated genes ($|\log_2 FC| \geq 1.0$, adjusted $p \leq 0.05$). Log2-transformed DEG counts were used for visualization.

### Supplemental

A.

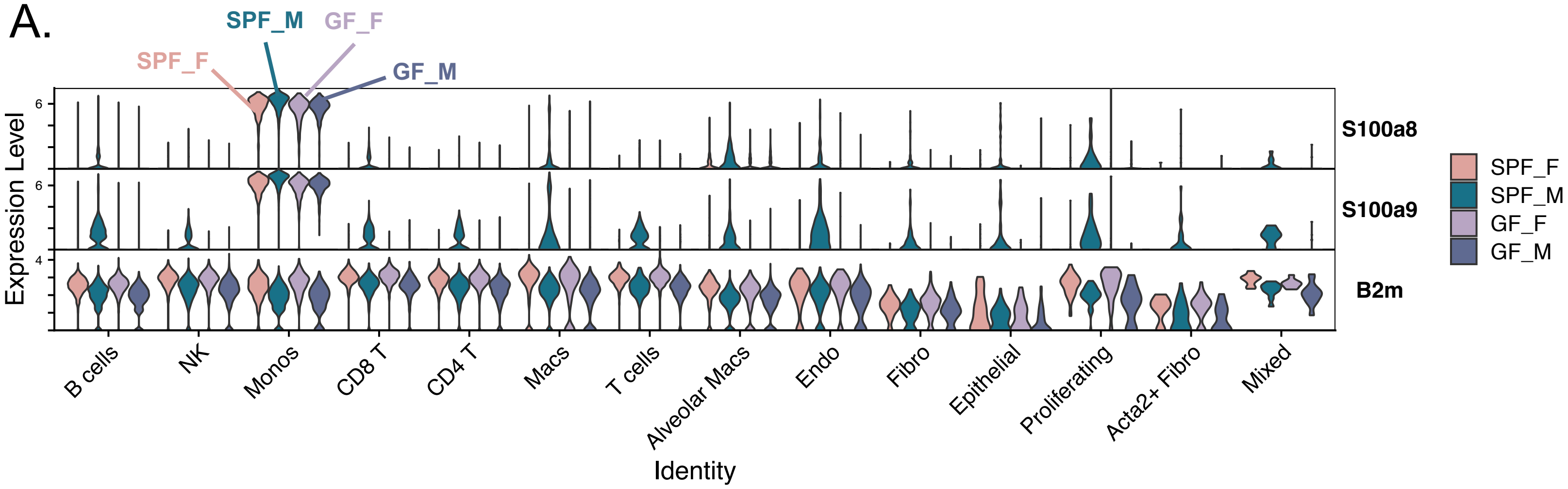


**Supplemental Fig 3. Additional data supporting Figure 4. A.** Violin plots illustrating expression of S100a8/9 and B2m across major cell types.
