## Supplemental Table 1 for "Single-Cell RNA Sequencing Reveals Commensal Microbes Amplify Sex-Specific Immune Programming in the Murine Lung"

| Supplemental Table 1. | | | | |
|---|---|---|---|---|
| **Fluorochrome** | **Antigen Target** | **Clone** | **Vendor** | **Catalog #** |
| Live-Dead Fixable Aqua | Live/Dead | Fixable Aqua Dead Cell Stain Kit | Invitrogen | L34965 |
| BV421 | CD45.2 (Ly5.2) | 104 | Biolegend | 109832 |
| BV605 | CD8 | 53-6.7 | Biolegend | 100744 |
| BV650 | CD45 | 30-F11 | Biolegend | 103151 |
| BV750 | CD19 | 6D5 | Biolegend | 115561 |
| Alexa Fluor 488 | Ly-6C | HK1.4 | Biolegend | 128021 |
| Spark Blue 550 | CD4 | GK1.5 | Biolegend | 100473 |
| PE | F4/80 | BM8 | Biolegend | 123109 |
| RB705 | Ly-6G | 1A8 | BD Pharmingen | 757648 |
| PerCP-Fire 806 | CD11b | M170 | Biolegend | 101293 |
| APC | NK1.1 | PK-136 | Biolegend | 108709 |
| APC-Fire 750 | CD3 | 145-2C11 | Biolegend | 100361 |
